# Multicenter HFpEF study identifies sex disparity linked with two discrete cardiac proteomic signatures

**DOI:** 10.64898/2025.12.18.695212

**Authors:** Aleksandra Binek, Johannes V. Janssens, Niveda Sundararaman, Yaqun Teng, Kim M. Mellor, Antonia J. A. Raaijmakers, Claire L. Curl, Lizhuo Ai, Lawrence S. C. Czer, Amy D Bradshaw, Kavita Sharma, Virginia S. Hahn, Kenneth S. Campbell, Michael R. Zile, David A. Kass, Eduardo Marban, Lea M.D. Delbridge, Jennifer E. Van Eyk

## Abstract

Heart failure with preserved ejection fraction (HFpEF) is epidemic, with an incidence exceeding that of heart failure with reduced ejection fraction (HFrEF). Sex differences in HFpEF phenotype have been observed, particularly linked to aging and metabolic comorbidities in women. Here, we report a multi-centre proteomic analysis of HFpEF and HFrEF cardiac samples. Using tissues and clinical data derived from multiple institutional biobanks, in-depth characterization of the proteome of ventricular samples was undertaken. Relative to HFrEF, the HFpEF cohort exhibited minimal proteome composition overlap and pronounced heterogeneity. A key finding of this investigation is that sex per se did not confer a distinctive proteomic HFpEF signature. Rather, two HFpEF proteomic profiles were identified, differing significantly in sex ratios. The identification of two Clusters revealed that HFpEF has two predominant proteomic signatures at the molecular level, marked by differences in the extent and nature of structural and contractile machinery and by local cellular and ECM communications. Upstream regulator analysis identified various molecular leads to be pursued in defining these two HFpEF profiles. Our findings offer specific opportunities for new exploration of therapeutic options to target the spectrum of HFpEF proteomic diversity and identify potential drug targeting prospects.

---

Heart failure with preserved ejection fraction (HFpEF) is epidemic, with an incidence exceeding that of heart failure with reduced ejection fraction (HFrEF).^1^ Sex differences in HFpEF phenotype have been observed, particularly linked to aging and metabolic comorbidities in women.^2^ While HFpEF is increasingly viewed as a complex syndrome involving a range of systemic disturbances, dysfunction of the heart underlies the characteristic pulmonary congestion and exercise intolerance.^1^ Molecular mechanisms remain mysterious, with studies of human HFpEF being constrained by limited access to heart tissue. Here, we report a multi-centre proteomic analysis of HFpEF and HFrEF cardiac samples, contrasting the two diseases, with particular focus on the molecular bases of HFpEF heterogeneity.

Collaborating authors shared patient biobank tissue samples and clinical data from each of their institutions, previously reported in relation to other investigations (see ‘Ethical Approval’, & references).^3.4.5.6^. The larger sample size facilitated the characterization of sex-specific signals and pathways. From each biobank (two HFrEF,^3,4^ two HFpEF^5,6^), ventricular samples from patients for whom the primary diagnosis was ‘heart failure’ were selected. To further refine sample clinical origins, criteria were applied to exclude additional diagnoses of amyloidosis, sarcoma, idiopathic and/or hypertrophic cardiomyopathy, evidence of ischemia and other confounding conditions. Samples (n=109) comprised 61 HFrEF (18 female, 43 male) and 48 HFpEF (29 female, 19 male)

Pooled clinical parameters (female and male, means ±SD) for the HFrEF group showed a low EF (%: 21.6±7.5 and 17.1±4.9), relatively young age (yrs., 53.4±13.2 and 50.8±11.3), and comorbid diabetes (44% and 47%). Pooled HFpEF data showed well-maintained EF (%: 65.3±6.1 and 66.3±5.8), an older age profile (yrs, 63.1±10.0 and 59.2±12.99), and increased diabetes (66% and 68%). Samples were identically processed at Cedars-Sinai Medical Center by mass spectrometry using ‘data-independent’ acquisition and analysed as described^7^. Principal Component Analysis (PCA) demonstrated minimal overlap of HFpEF and HFrEF groups, and relatively high HFpEF proteomic heterogeneity (Figure 1A).

**Figure 1.**
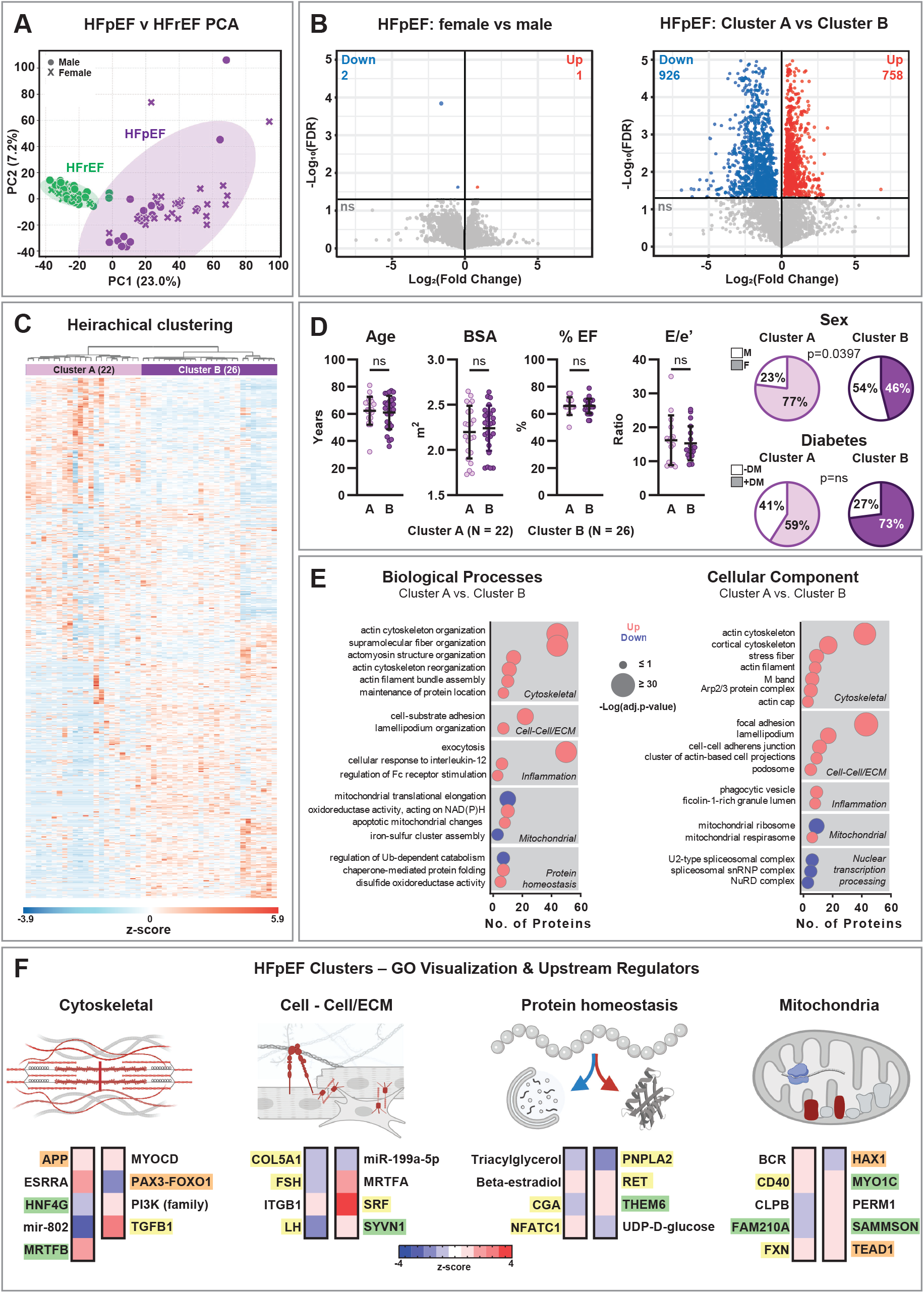
Proteomic analysis of cardiac samples from HFrEF and HFpEF patients. Samples were processed for mass spectrometry and proteomics performed using ‘data-independent acquisition’ methods and analysed as previously detailed.^7^ All raw MS data were searched together against *Homo sapiens* Uniprot database (https://www.uniprot.org/help/homo_sapiens) and quantified using DIA-NN (Data-Independent Acquisition by Neural Networks, V1.8.1). Data processing steps included a cohort normalization (protein counts), application of 50% missingness filter, imputation by 1/5 of minimum protein value. **(A). Principal Component Analysis (PCA) of HFrEF & HFpEF patient group proteomics**. Dimensional reduction of the mean-scaled protein abundance data was performed using PCA. PC1 (x-axis) and PC2 (y-axis) are displayed as scatterplots for HFpEF (purple) and HFrEF (green) overlaid with a 95% co-variance ellipse. Data points are designated female or male. **(B). Volcano plots depicting sex and HFpEF Cluster proteomic comparisons**. Sex (left, female vs male) and Cluster (right, Cluster A vs. Cluster B) comparisons are shown. Data are shown as Benjamini-Hochberg (BH) adjusted p-values (-log10, y axis) vs fold change (log2fold change, x-axis) visualized in ‘volcano’ plot form. Student’s t-test, with BH adjustment and ‘False Discovery Rate’ (FDR) set at p=0.05 (ie 5%). Significantly increased (red) and decreased (blue) protein abundance fold-change shown, with non-significant proteins as grey. **(C). Heatmap and dendrogram depicting hierarchically clustered HFpEF proteomic data**. An unsupervised hierarchical clustering approach was applied to the mean-scaled (z-scored) protein abundance data (‘MetaboAnalyst’ V6). Parameters settings (default) included ‘Euclidian distance’ and ‘Ward’s Method’ linkage. The dendrogram depicts two high level Clusters of different sample sizes: Cluster A (n=22) and Cluster B (n=26). The z-score scale indicates higher (red) and lower (blue) protein abundance. **(D). Patient clinical metadata and proteomic determined HFpEF Cluster comparisons**. The clinical parameters age (years), body surface area (BSA, m^2^), cardiac ejection fraction (EF, %) and diastolic function (E/e’, ratio) are presented as scatter plots for patients comprising Cluster A (n=22) and Cluster B (n=26) presented as mean±std dev (Mann-Whitney U-test, all p=ns). Pie-charts show the sex proportions for each Cluster (% female: A=77, B=46; p=0.0397 Fisher Exact test) and the proportion of patients diagnosed as diabetic comprising each Cluster (% diabetes: A=59, B=73; p=ns, Fisher Exact test). Statistical analyses performed in Prism Graph Pad (V10.6.1). **(E). HFpEF patient proteomic Cluster comparisons using ontological protein classifications**. The Cluster-identified data set was analysed using a linear (limma) model, specifying ‘batch’ for adjustment as a single covariate (BH adjustment as FDR ≤ 5%) to minimize any possible effect of sample processing in multiple groups. The ranked GO output from the adjusted linear model was further processed using the Protein Interaction Network Extractor (PINE) tool coupled with GO enrichment exploration with the ClueGO utility. Refer [https://github.com/csmc-vaneykjlab/pine] and [Cytoscape https://apps.cytoscape.org/apps/cluego]. With PINE, increased (red) and decreased (blue) regulated terms were screened with high stringency (Bonferroni adjustment as FDR ≤ 1%). GO term redundancy was reduced by grouping terms based on the similarity of associated proteins using ClueGO (kappa score @ 0.5). The most significant GO term within each ClueGO group of terms was designated the ‘Leading term’. A total 37 Leading terms were identified in the Biological Process (BP, n=18) or Cellular Compartment (CC, n=19) GO categories. The ClueGO groups are listed in the Panel, organized into BP and CC Theme groups based on Leading term descriptor information. Each of the ClueGO Leading terms are depicted as symbols with attributes which describe Cluster A vs Cluster B term differences. Symbol colour denotes abundance difference (red higher & blue lower), symbol diameter denotes significance of difference (-log_10_ transformed, p-value, 5%FDR), and symbol position on the x-axis denotes protein count for each Leading term. **(F). Identifying upstream regulators of potential pathologic importance in HFpEF Clusters**. Cluster-based upstream regulator analysis was performed in Ingenuity Pathway Analysis (IPA V1.23.1) software using fold-changes and adjusted p-values (p<0.05) for differentially expressed proteins identified in the limma linear model analysis. In total, 48 significant endogenous upstream regulators were identified (FDR 5%). Of these, 28 regulators were predicted to be activated (Z-score ≥ 2) and 20 inhibited (Z-score ≤ -2). Overlap between proteins linked with each predicted upstream regulator and proteins linked with each GO Leading term (Panel E) was assessed (Fisher Exact test, SciPy V1.16.13). Of the 48 upstream regulators, 35 were significantly associated with Leading GO terms (p<0.05). Upstream regulators (activated or inhibited) were assigned to Theme groups based on strongest overlap (Odds Ratio) of protein networks between each upstream regulator and each GO Leading term. The BP and CC themes with the largest count of assigned upstream regulators (Cell-Cell /ECM (n=8), Cytoskeletal (n=9), Mitochondrial (n=10), Protein homeostasis (n=8)) are <u>depicted</u> as motifs illustrating sub-cellular and trans-cellular protein localization sites. Red/Blue colour is used to identify loci of potential HFpEF Cluster phenotype differences. Upstream regulators are shown below each theme motif as ‘heat strips’ (red = activated, blue = inhibited) where the colour intensity represents magnitude of the IPA activation z-score. The ‘druggable’ characteristics of upstream regulators were evaluated using the ‘Biofinder’ Chemical Abstract Service (CAS, American Chemical Society) and literature search. Druggable regulators have been described using colour-coding. For <u>‘promising</u> targets (4, orange highlight) investigations ongoing have so far not exposed cardiac risk. Other <u>potential</u> targets (7, green) have been pursued but not progressed to human study to date. Regulators already linked with cardiovascular risk or cardiotoxic actions are shown as precluded (11, yellow). The non-coloured regulators are those for which ambiguous, variable or no findings are reported. [Figure created in BioRender. Mellor, K. N28ZSA7HP].

Next a sex-specific analysis was performed for the HFpEF group. Only a few proteins were identified as significantly different between the sexes (reassuringly including a sex-chromosome related protein, Figure 1B, left). Thus, sex, as a binary categorical variable, did not constitute a cardiac proteomic signature in this set of HFpEF samples. In contrast, ‘unsupervised’ clustering of the same data, generated a hierarchical proteomic map composed of two HFpEF clusters: A (n=22) and B (n=26) comprised of both sexes (Figure 1C). In a cluster-specific analysis, 38.2% of the observable proteome was altered (Figure 1B, right). Of 4411 proteins quantified, 1684 proteins exhibited significant differential abundance (758 increased & 926 decreased). Comparison of patient metadata for Clusters A and B (Figure 1D) revealed no differences in age, body surface area (BSA), ejection fraction (EF) or diastolic function (E/e’, which showed impaired diastolic relaxation). Although clinical characteristics were comparable, major underlying Cluster proteomic identities were discerned (Figure 1D). When comparing Clusters A and B, the percentages with diabetes were not different, but Cluster A had a higher percentage of women (% female: A=77 *vs*. B=46, Figure 1D, right).

To probe pathogenic characteristics differentiating Clusters A and B, we used a linear covariate analysis with a gene ontology (GO) investigation (Figure 1E). This model refined the count of differentially abundant proteins to 346 (187 increased,159 decreased in A vs. B). Of 110 significant GO terms, 69 represented Biological Processes (BP) and 41 Cellular Components (CC). ClueGO utility analysis collapsed similar terms under Leading terms (BP, n=18; CC=19; total=37), which were grouped in Themes shown in Figure 1E.

Cluster A (predominantly female demographic) exhibited a marked elevation in abundance of sarcomere and cytoskeletal associated proteins (‘<u>Cytoskeletal</u>’) compared with Cluster B (balanced sex representation), with upregulation of proteins involved in extracellular matrix (ECM) and adjacent cell interactions (‘<u>Cell-Cell/ECM’</u>). These cell and matrix features have been implicated in modulating diastolic stiffness. Similarly, the Clusters differed in the abundance of chaperones involved in protein folding and mediators of misfolded protein degradation (‘<u>Protein homeostasis’</u>) - processes linked to HFpEF.^1^ Alterations in mitochondrial proteins indicated modified electron transport chain function involving complexes I and III, and decreased mito-ribosomal abundance (‘<u>Mitochondria</u>’). Cluster-specific differences in proteins linked with inflammation and nuclear transcription were also evident. Notably absent in Cluster comparison was evidence of protein abundance indicative of major cellular metabolic disturbance and autophagic processing. In studies where HFpEF samples are compared with ‘control’ or other disease states this is a prominent finding.^1,6-9^ An inference is that both HFpEF Clusters included similar protein dysregulation relating to these pathologies.

To interrogate pathways involved in driving HFpEF proteome Cluster differences (Figure 1F), we performed Ingenuity Pathway Analysis (IPA) upstream regulator analysis. Upstream regulators (activated or inhibited, n=35) were assigned to each Theme group based on strongest overlap (Odds Ratio) of protein networks between each regulator and the GO Leading term. Upstream regulator data, shown as ‘heat-strips’ adjacent to illustrative Theme motifs (Figure 1F), highlighted 9 transcription factors (ESRRA, HNF4G, MRTFB, MYOCD, MRTFA, SRF, NFATC1, CLPB, TEAD1), many of which relate to Cytoskeletal and ECM structure and function. Also identified were two microRNAs (mir-802, miR-199a-5p), both previously reported in cardiac tissue. Several endocrine mediators were recognized (beta estradiol, FSH, LH), possibly underlying some of the sex differences between HFpEF Clusters. Well-characterized signalling family molecules (PI3K, TGFB1) and metabolic species (UDP-glucose, triacylglycerol) were represented. The ‘druggable’ characteristics of upstream regulators were evaluated, and colour highlighted to show ‘promising’ (orange), ‘potential’ (green) or already ‘precluded’ (yellow) candidates (Figure 1F legend).

Whilst analysis of multiple tissue sets provides the opportunity to expand the scope and power of research (especially in a field where tissue samples are scarce), there are inherent limitations. Although tissue and clinical data were derived from geographically distributed sources, and anatomic locations of ventricular samples were not uniform, inclusion of rigorous standardized proteomic methodologies mitigated data variability.

To summarize, tissue sets derived from multiple institutional biobanks enabled in-depth characterization of the proteome of ventricular samples from a defined group of HFpEF patients. Relative to HFrEF, the HFpEF cohort exhibited minimal proteome composition overlap and pronounced heterogeneity. A key finding of this investigation is that sex *per se* did not confer a distinctive proteomic HFpEF signature. Rather, two HFpEF proteomic profiles were identified, differing significantly in sex ratios. The identification of two Clusters revealed that HFpEF has two predominant proteomic signatures at the molecular level, marked by differences in the extent and nature of structural and contractile machinery and by local cellular and ECM communications. Upstream regulator analysis identified various molecular leads to be pursued in defining these two HFpEF profiles. Our findings offer specific opportunities for new exploration of therapeutic options to target the spectrum of HFpEF proteomic diversity and identify potential drug targeting prospects.

## Acknowledgements

**JVJ** gratefully acknowledges financial support from the Fulbright U.S. Scholar Program, which is sponsored by the U.S. Department of State and Australian-American Fulbright Commission.

## Declarations

### Data Availability

Data and materials supporting the results or analyses presented in this paper are available on reasonable request to the corresponding author. Mass Spectrometry data are available in Proteomics Identifications Database PRIDE (identifier PXD045677, https://www.ebi.ac.uk/pride/archive).

## Ethical Approval

All samples were collected and stored in accordance with the Guidelines for Human Tissue Repository of the NIH (National Heart, Lung and Blood Institute). All patients diagnosed with reference to established protocols/guideline statements. All sample collection protocols occurred with relevant Institutional Review Board (IRB) approvals.

Sample information and primary source data reported (by PMID) summarized: Medical University of South Caolina, Charleston, SC, USA.

IRB: 00054823 (PMID:25637629)^5^. LV anterior free wall sub-epicardial biopsies. Cedars Sinai Medical Center, Los Angeles, California, USA.

IRB:11910 (PMID:31292486)^4^. Ventricular apical transmural cylindrical piece of myocardium. Johns Hopkins University, Baltimore, MD, USA.

IRB: on request (PMID:33118835)^6^. Right ventricular septal endomyocardial biopsies. University of Kentucky, Lexington, KY, USA.

IRB: 08-0338-F2L (PMID:29081749)^3^. Pre transplant, explanted heart ventricular endomyocardium.

